# Speech prosody supports speaker selection and auditory stream segregation in a multi-talker situation

**DOI:** 10.1101/2022.04.12.487484

**Authors:** Petra Kovács, Brigitta Tóth, Ferenc Honbolygó, Orsolya Szalárdy, Anna Kohári, Katalin Mády, Lilla Magyari, István Winkler

## Abstract

To process speech in a multi-talker environment, listeners need to segregate the mixture of incoming speech streams and focus their attention on one of them. Potentially, speech prosody could aid the segregation of different speakers, the selection of the desired speech stream, and detecting targets within the attended stream. For testing these issues, we recorded behavioral responses and extracted event-related potentials and functional brain networks from electroencephalographic signals recorded while participants listened to two concurrent speech streams, performing a lexical detection and a recognition memory task in parallel. Prosody manipulation was applied to the attended speech stream in one group of participants and to the ignored speech stream in another group. Naturally recorded speech stimuli were either intact, synthetically F0-flattened, or suppressed by the speaker. Results show that prosody – especially the parsing cues mediated by speech rate – facilitates stream selection, while playing a smaller role in auditory stream segmentation and target detection.

## 1. Introduction

The human brain typically solves the problem of processing speech in a multi-talker environment seemingly effortlessly (Cherry, 1953). This goal is reached by separating the mixed acoustic input into distinct, coherent streams of sound/speech (termed auditory stream segregation; Bregman, 1994) and selects one of them for in-depth processing (selective attention; Broadbent, 1952). Both of these functions may involve higher cognitive operations, necessitating cooperation between several brain areas (Hill & Miller, 2009; Saur et al., 2008, 2010). Although prosody is known to aid speech comprehension (Carlson, 2009; Kjelgaard and Speer; 1999, LaCroix et al., 2020; Roncaglia-Denissen et al., 2013; Sheppard et al., 2017), its role in processing speech in multi-talker environments is less well known. The present study aims to address this issue.

Prosody can be defined as the melody of speech, which carries linguistic information and information about the speaker’s mental state. Prosody constitutes many features of speech, including intonation (which is the variation of the fundamental frequency, or F0 of speech), stress, rhythm, timing, and pauses (Myers et al., 2019; Sammler et al., 2015). Myers and colleagues (2019) noted that prosody is an important feature of speech from the viewpoint of comprehension: the different features of prosody all assist the segmentation and the parsing of speech input into meaningful units at different levels of processing (i.e., sentence, phrase, word). Prosody is processed automatically and without conscious effort by the listeners (Frazier et al., 2006). Although languages differ in the way they implement prosody, each language uses some form of prosodic grouping and prominence marking (e.g., stress, edge-prominence, etc.) Prosodic grouping may also be used to provide the basic structure of utterances, a scaffolding that allows listeners to hold an auditory sequence in memory. This suggestion is supported by empirical findings showing that sentences heard with typical prosody are better understood compared to those with atypical prosody (Carlson, 2009; Kjelgaard and Speer; 1999, LaCroix et al., 2020; Roncaglia-Denissen et al., 2013; Sheppard et al., 2017). It is possible that sentences with typical prosody allow a more efficient use of cognitive resources, because they focus the listener’s attention to the important elements, and prosodic boundaries divide utterances into smaller units to be parsed. Within the brain, prosody is assumed to be predominantly processed in the right hemisphere (Ross & Monnot, 2008; Witteman et al., 2011). Sammler and colleagues (2015) suggested a dual-route model for processing prosodic information: a dorsal stream including the right premotor cortex and inferior frontal gyrus and a ventral stream including the right posterior and anterior superior temporal sulcus.

Multiple prosodic features may play a role in directing attention. Firstly, changes in fundamental frequency (F0) and pauses can signal boundaries between speech units. Secondly, speech rate can affect comprehension and memory processes. In a multi-talker situation, Szalárdy and colleagues (2020) showed that fast speech rates result in a trade-off between two concurrent tasks (target detection and memory encoding), with the performance in one or both decreasing at fast speech rates. This suggests temporally limited availability of some processing capacity required by both tasks. Thus prosody is expected to play a role in the selective processing of the designated speech stream in a multi-talker situation through guiding attention and delineating processing units.

Spectral analysis of electro-/magnetoencephalographic (EEG/MEG) data recorded during speech processing led to important observations about the neural activity underlying attentional processes related to speech comprehension. It has been shown that the phase of low-frequency delta/theta activity (<8 Hz) synchronizes with the temporal structure of speech (termed neural tracking; Luo & Poeppel, 2007). In the case of two concurrent speakers, phase-locking occurs for both speech streams, with the attended stream being preferentially represented in the posterior auditory cortex (Ding & Simon, 2012) and results in stronger synchronization (Rimmele et al., 2015). Teoh, Cappelloni and Lalor (2019) recently showed that prosody tracking within the EEG signal can be separated from the tracking of other speech cues: neural oscillations in the delta band were shown to be entrained to relative pitch for intact speech, while the prosody of vocoded speech was not tracked. Similarly, Rimmele and colleagues (2015) found that synchronization deteriorated when the acoustic quality of the attended stream was degraded. In sum, neural speech tracking is enhanced by attention, but this enhancement is dependent on the quality of the acoustic signal.

As for selective attention effects appearing in EEG signals, increases in alpha power (∼10 Hz) have been linked to the selective inhibition of the distracting speech stream (Strauß, Wöstmann & Obleser, 2014). This mechanism represents top-down processes, because alpha power enhancement occurs later than low-frequency neural tracking, and originates from higher order brain areas such as the fronto-parietal attention network (Wöstmann et al., 2016; Fiedler et al., 2019).

One way to characterize the brain networks involved in the above mentioned processes is to calculate functional connectivity (FC) between regions (e.g., Stam et al., 2007). For instance, Obleser et al. (2007) showed that understanding acoustically degraded but semantically predictable speech is accompanied by enhanced FC between cortical regions outside the auditory cortex, including prefrontal, inferior frontal, and posterior cingulate areas.

The present study addressed the question to what degree prosodic characteristics of the attended and ignored speech stream promote speaker segregation and/or improve the selection of the to-be-attended stream. To this end, participants were presented with two concurrent speech streams, the prosody of which was manipulated on three levels (intact, synthetically flattened F0, and naturally suppressed – i.e., spoken in a monotonous voice). The to-be-attended speech stream was manipulated for one group of participants, while the to-be-ignored stream was manipulated for the other group. Listeners were instructed to detect numeral words in the attended stream.

Three kinds of data were collected and analyzed in the current study: behavioral measures, event related brain potentials (ERP), and functional connectivity (FC). All three carry information about different aspects of speech processing in a multi-talker situation. Behavioral tasks (numeral detection and recognition memory tests) ensure that listeners pay attention to the appropriate speech stream during the experiment, and they help us objectively evaluate how difficult it was to attend a given speech stream and suppress distractors. In comparison, ERP data show the momentary changes in neural activity during the numeral detection task. Two successive ERP components, the N2b and the P3b are typically elicited for target auditory stimuli (Näätänen et al., 1982; Polich, 2007; Polich & Herbst, 2000; Ritter et al., 1983). The N2b is a negative component with a centro-parietal scalp distribution which usually peaks at around 200 ms from stimulus onset, and appears after a detected target event. It has been associated with stimulus classification (Näätänen, 1990; Ritter et al., 1979), and studies have shown that its amplitude is modulated by selective attention (Michie, 1984) and stream segregation (Szalárdy et al., 2013). For target events, the N2b is usually followed by the P3b, which is a positive ERP component with a parietally dominant scalp distribution and maximum amplitude at around 300–400 ms from stimulus onset (Conroy & Polich, 2007). P3b has been associated with context updating, categorization, and later evaluation of the target stimulus, and interaction between selective attentional processes and working memory (Polich & Herbst, 2000). Compared to behavioral data, ERPs generally carry additional information, such as that healthy elderly often perform at the same level as young participants in an auditory figure-ground segregation task, while their ERP responses during the completion of the task show compensatory efforts (Boncz et al., 2021). Thus, whereas participants may improve their behavioral performance through conscious effort, the compensatory processes modulate event-related neural activity, making the two measures more informative together. Lastly, we extracted EEG functional connectivity data. The difference between the ERP and the FC approach lies in the fact that unlike ERPs, FC is not dominated by processes related to some specific stimulus or response event. Instead, it reflects processes on a longer time scale, including preparatory and context-related processes, and thus can show the long-term brain activity changes subserving adaptation towards successful task performance. Further, in terms of brain topography, FC reflects interactions between various regions of the neocortex. In sum, we expected that each of these measures were affected by prosody, and that they provide information about different aspects of how participants cope with the experimental situation.

The following hypotheses were formed. I) The deterioration of prosodic cues will adversely affect the selection of the designated stream and/or the target within it, but not as much auditory stream segregation itself, because there are many acoustic differences between speakers, of which prosodic differences constitute only a part. This hypothesis would be confirmed by larger behavioral and neurophysiological effects for the attended stream manipulated group than the ignored stream manipulated group. Regarding functional connectivity, we expect that slow oscillations (delta, theta) would link the cortical regions involved in selective attention. When it is more difficult to selectively attend to the target stream, stronger coupling is expected between the brain regions related to focusing on the attended stream. In parallel, finding increased alpha-band activation would suggest an increased effort to suppress the unattended stream.

II) Stimuli created by the speaker “naturally” suppressing prosody will have less effect on the listeners’ performance and brain activity than those created by the synthetic prosody suppression method, as the former retains some F0 variation. This hypothesis would be confirmed by larger behavioral effects for the synthetic than for the natural prosody suppression conditions. Further, the alpha brain networks should also differentiate between the two prosody manipulations on the attended stream, showing differences in the effort needed to suppress the unattended stream (see above).

## 2. Materials and methods

### 2.1. Participants

Fifty healthy young adults, all native speakers of Hungarian were randomly assigned to one of two experimental groups: “attended stream manipulated” (N = 25; 18 female; mean age: 21.2 years, SD = 1.44; 24 right-handed) or “ignored stream manipulated” (N = 25; 19 female; mean age: 21.92 years, SD = 2.45; 21 right-handed). No history of psychiatric or neurological symptoms was reported by any of the participants. All participants had pure-tone thresholds ranging from 250 Hz to 4 kHz: <25 dB and <10 dB difference between the ears, as determined by audiometry. Informed consent was signed by all participants after the aims and methods of the study were explained to them. The study was conducted in full accordance with the World Medical Association Helsinki Declaration and all applicable national laws; it was approved by the institutional review board, the United Ethical Review Committee for Research in Psychology (EPKEB). Listeners received modest financial compensation for their participation.

A participant’s data would have been excluded if less than 90 artifact-free trials could be extracted for at least one condition, but no participant violated this criterion.

### 2.2. Stimuli

In each stimulus block, participants were presented with two concurrent speech streams of ca. 6 minutes duration each (mean duration: 352.15 s, SD = 9.34). The stimuli were news articles, which were reviewed to ensure correct grammar, natural text flow, and no garden path sentences. The information in the articles was obtained from Hungarian news websites, and the articles contained emotionally neutral, not too well-known facts. Twelve–twelve articles were delivered in the experimental session by two male native Hungarian speakers (professional actors) with one article from one and another from the other speaker, concurrently (24 articles in total). The soundtracks were recorded at 48 kHz with 32-bit resolution and edited by a professional radio technician. They were presented by Matlab R2014a software (Mathworks Inc.) on an Intel Core i5 PC with ESI Julia 24-bit 192 kHz sound card connected to Mackie MR5 mk3 Powered Studio Monitor loudspeakers. The recording site was the same room where later the experiment took place. During the experiment, sounds were delivered from the approximate location of the speaker of the recording (i.e., the loudspeakers were mounted at approximately the same location as the actors’ heads). A constant location was assigned to each speaker: one on the left, the other on the right side of the listener (Figure 1A).

**Figure 1.**
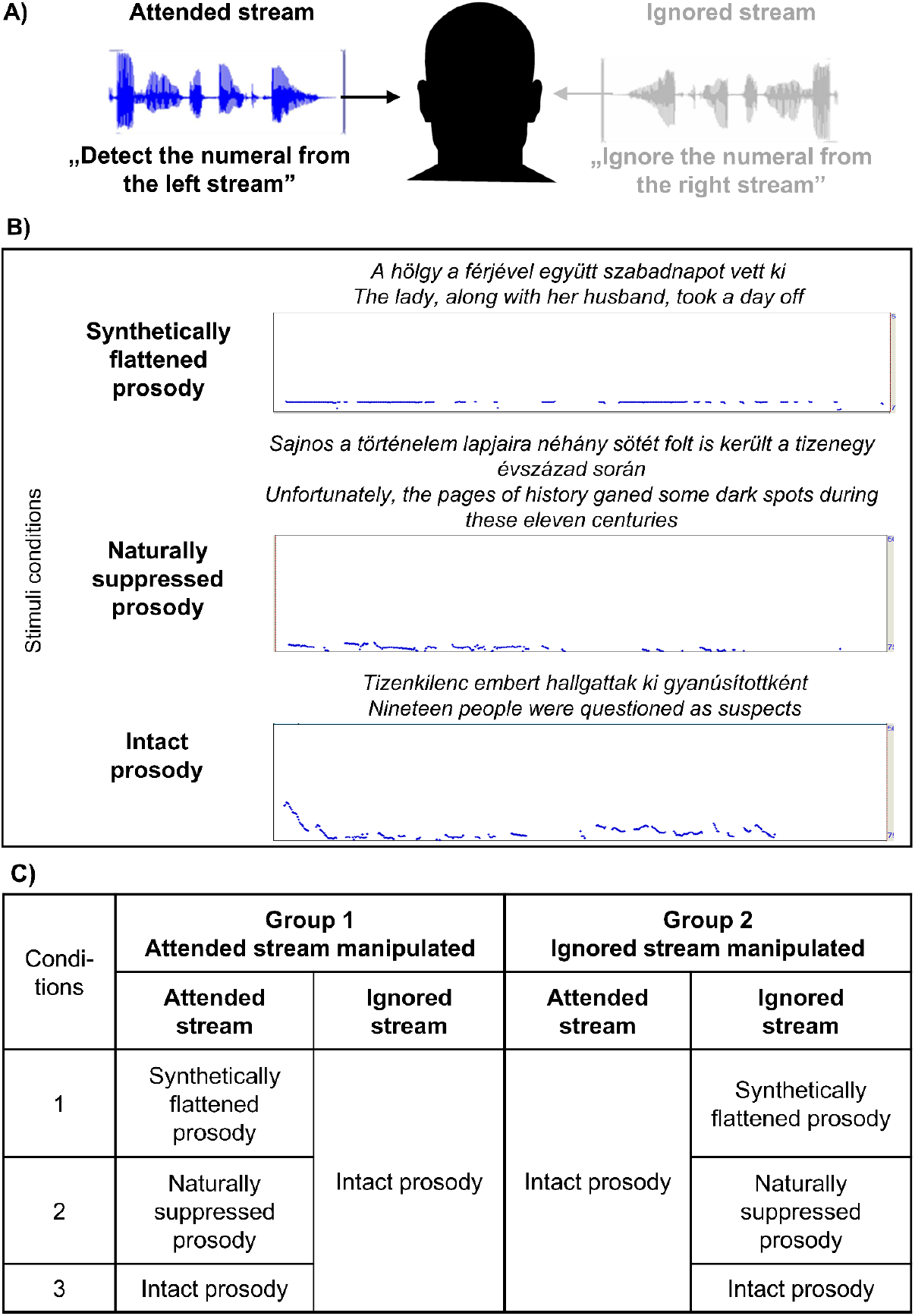
Experimental design. A) Participants listened to two concurrent acoustic streams, detecting numerals by pressing a response key for the stream delivered from the left side. B) Segments were either intact speech, speech with synthetically flattened prosody, or speech with naturally suppressed prosody. The original Hungarian example sentences with their speech envelopes are shown with their English translation. C) Table of the conditions for the attended stream and the ignored stream manipulated groups.

Three types of stimuli were created from the 24 articles delivered for the manipulated side/speaker (four for each stimulus type), while for the unmanipulated side/actor, intact speech was presented (see Figure 1B). The same articles were delivered to both groups. The manipulated articles were substituted by their intact version when they fell on the unmanipulated side/actor of the group.

1. <u>Intact speech:</u> no manipulation was employed on the speech signal.
2. <u>Speech with synthetically flattened prosody:</u> We followed the method used by Keitel et al. (2013). To create the sentences with the flattened intonation, we removed the intonation contours and replaced them with the average of the mean F0 of the whole block of sentences using the Praat software (Boersma & Weenink, 2022): the pitch contour of the sentences was extracted and segmented into pitch points at a rate of 100 Hz, after which the pitch points were removed and replaced with a new pitch contour with the average frequency of the respective recording using PSOLA (pitch synchronous overlap and add) resynthesis;
3. <u>Speech with naturally suppressed prosody:</u> The actor was asked to deliver the sentences in a monotonous way, “as if it was delivered by a machine”.

Prosodic differences between the three stimulus types were assessed separately for speech rate and pitch variation (Table 1). Speech rate was assessed on two randomly selected sample recordings from each condition (one for each speaker) by the lmerTest package in R (Kuznetsova et al., 2017; R Core Team, 2020) using linear mixed-effects models. Sample recordings were annotated in Praat (Boersma & Weenink, 2022) with regard to sentence boundaries, including pauses. Because the current synthetic manipulation does not affect speech rate, this analysis only included the synthetically and the naturally manipulated stimulus types. In the mixed-effects model, a nested random effect was included: sentences nested within speakers. The results showed that speech type had a significant effect on speech rate (*β* = −0.456, *t* = −11.44, p < 0.001, marginal R^2^ = 0.20, conditional R^2^ = 0.68): speech was faster in the naturally prosody-suppressed than in the synthetically flattened speech type. Note that the acceleration of speech rate could not only stem from faster articulation, but also from shorter or missing pause intervals, showing a less structured prosodic phrasing in general.

**Table 1.** Prosody-related acoustic parameters of two sample recordings from each stimuli condition: median (Md), standard deviation (SD), interquartile range (IQR) and range of pitch, and mean (M) and SD of speech rate.

|  |  | Synthetically flattened speech | Naturally prosody-suppressed speech | Intact speech |
| --- | --- | --- | --- | --- |
| Pitch (Hz) | Md | 147.41 | 117.08 | 131.19 |
|  | SD | 3.88 | 14.63 | 35.60 |
|  | IQR | 0.67 | 21.16 | 61.87 |
|  | Range | 180.45 | 198.25 | 224.00 |
| Speech rate (syllables/s) | M | 4.81 | 5.26 | 4.81 |
|  | SD | 0.45 | 0.36 | 0.45 |

Pitch range was assessed on the whole set of recordings for all three conditions with a Friedman ANOVA test. Mean pitch range was significantly affected by speech type (X _F_^2^(2) = 14.25; p < 0.001), with synthetically flattened speech (M Rank = 1.13) having a significantly smaller pitch range than intact speech (M Rank = 3.0; p_bonf_ < 0.01) according to Conover’s post-hoc comparison. In sum, whereas the synthetic manipulation only affected variation in the fundamental frequency, speakers instructed to speak monotonously reduced pitch variation to a lesser degree, but also increased their speech rate.

Each article contained 42–54 numerals (M = 49.79, SD = 3.16), which served as targets in the attended stream and as distractors in the ignored stream. Additionally, articles in the ignored but not the attended stream contained 19–21 syntactic violations (M = 20.08, SD = 0.90) included for the purpose of determining whether or not the to-be-ignored stream was not (at least intermittently) attended by the participants. Consciously processed as opposed to ignored syntactic violations elicit the ELAN, LAN/N400, and/or P600/SPS ERP component (Friederici, 2002), as was also shown by Szalárdy and colleagues (2018), who used a setup closely similar to that of the current study.

### 2.3. Procedure

Figure 1 illustrates the experimental design. The two groups differed only in whether the prosody manipulations were applied to the attended or the ignored stream. The other stream always consisted of intact speech (Figure 1C).

The experiment was conducted in an acoustically attenuated and electrically shielded room. In order to decrease motor activity and eye blinks during the experiment, participants were instructed to focus on a fixation cross at the center of a 19” monitor placed directly in front of them. Participants were simultaneously presented with two speech streams from two loudspeakers, one positioned at 30° left from the frontal midline, while the other symmetrically to the right, both placed at a distance of ca. 2 m from the participant.

Participants performed two concurrent tasks. The first task was to detect numerals appearing within the article heard in the left (attended) stream, ignoring the distractor numerals in the right stream. Participants were instructed to respond to targets as quickly as possible with the press of a response key. Note that there was a confound between location and actor: participants always attended the same actor (the one whose voice was delivered by the left loudspeaker). After each stimulus block, a recognition memory test was performed. Five multiple-choice questions were asked with 4 possible answers each. The questions were related to the information in the article recently delivered in the attended stream. After the question and the 4 possible answers were read by the experimenter, the participant indicated his/her answer verbally. Participants were then asked to indicate whether they knew the answer from a different source than the recently presented text, in which case the question/answer was excluded from analysis.

Each participant received four stimulus blocks for each of the three stimulus conditions, resulting in 12 blocks/participant. The 6th block was followed by a mandatory break, with occasional shorter breaks between blocks in case the participant requested one. The six stimulus blocks of the first half of the session consisted of 2 blocks for each condition delivered in pseudorandom order with the constraint that consecutive stimulus blocks of the same condition were avoided. The order of the conditions was then reversed in the second half of the session. The order of the manipulated articles was randomized separately within each condition, as well as for the set of 12 concurrent intact-speech articles. No article was delivered twice during a session.

### 2.4. Data analysis

#### 2.4.1. Behavioral responses

In determining the temporal window of hits, we followed the method applied in our previous studies (e.g., Tóth et al., 2019). Responses were initially extracted from between 0 and 5000 ms from the onset of target events (numerals spoken in the attended stream). Responses were regarded as hits if they were longer than 5% and shorter than 95% of all responses of all individuals, resulting in an effective time window of 405–1689 ms.

Button presses following distractor numerals in the ignored stream (within the same temporal window as used for hits) were used to calculate the distractor effect, while false alarms were defined as responses to any non-target and non-distractor event. The importance of distinguishing between FAs and the distractor effect lies in the multi-talker design of the present study, due to which responses induced by numerals in the *to-be-ignored stream* were of specific interest. This is because distractor-induced responses imply that the two streams were confused by the listener; i.e., they are a proxy for the failure of stream segregation and/or selection. Lastly, d’ (Green & Swets, 1988) was calculated from the z-transforms of these variables as d’ = z(HITS) - z(DISTRACTOR EFFECT). Note that as a consequence, the d’ values show how well participants distinguished the two streams, not how well they distinguished numerals from non-numerals.

Recognition memory performance was separately calculated for each participant and condition. Items (questions) with an overall correct response rate above 95% or below 30% (collapsed across participants from both groups; 25% correct represents the chance level) were excluded from further analyses. No items needed to be discarded due to >95% correct response rate; 7 items with <30% correct response rate were discarded from the total of 120 items (12 × 5 = 60 item/group). Note that due to the random assignment of the intact-speech articles across the different conditions, the same article (and thus the same questions) could have appeared in different conditions for different participants (but each only once per participant). Recognition memory performance was then calculated separately for each listener (with reported knowledge from a different source taken into account) as the percentage of correct responses pooled across the stimulus blocks, separately for the three experimental conditions.

#### 2.4.2. EEG recording and preprocessing

EEG was recorded by a BrainAmp DC 64-channel EEG system with actiCAP active electrodes. Recordings commenced a few seconds before the onset of the article pair and concluded a few seconds after the shorter article ended. 62 electrodes were placed according to the International 10/20 system with FCz serving as the reference electrode. The sampling rate was 1 kHz, with a 100 Hz online low-pass filter applied. For monitoring eye movements, two electrodes were placed lateral to the outer canthi of the eyes. Impedances were kept below 15 kΩ during the recording.

EEG data was preprocessed with the EEGlab 11.0.3.1.b toolbox (Delorme, Sejnowski & Makeig, 2007) implemented in Matlab 2018b. Using a finite impulse response (FIR) filter (Kaiser windowed, Kaiser β=5.65, filter length 4530 points), an off-line band-pass filter was applied between 0.5 and 80 Hz. A maximum of two malfunctioning EEG channels were allowed for each participant.Malfunctioning channels were interpolated using EEGlab’s default spline interpolation algorithm. For artifact removal, the Infomax algorithm of Independent Component Analysis (ICA) was employed (Delorme, Sejnowski & Makeig, 2007). The topographical distribution and frequency contents of the ICA components were visually inspected, and the components constituting oculomotor artifacts were removed.

#### 2.4.3. ERP data analysis

From the continuous EEG recordings, epochs were extracted between -200 and +2400 ms relative to the onset of target and distractor numerals, as well as the syntactic violations appearing in the to-be-ignored articles. Baseline correction was applied by averaging the voltage in the -200–0 ms time window. Epochs exceeding the threshold of +/-100 µV change throughout the whole epoch measured at any electrode were rejected. The remaining epochs were averaged separately for each condition and participant. The mean number of items with accepted responses in the attended stream manipulated group were: 155.307 (77.7% of all trials, range: 79–203) target numerals, 186.72 (93.3%, range: 117–204) distractor numerals, and 78.64 (95.4%, range: 58–101) syntactic violations; in the ignored stream manipulated group: 159.427 (79.7%, range: 85–200) target numerals, 190.28 (95.1%, range: 137–254) distractor numerals, and 78.64 (98.3%, range: 58–101) syntactic violations. As expected (Szalárdy et al., 2018), target numerals elicited the N2b and P3b components, both with a maximum at parietal electrodes (see Supplementary Figure 1). Therefore, the time window for amplitude measurements was determined by setting intervals around the grand-average peak at the Pz electrode. As a result, the N2b amplitude was averaged from the 150–280 ms interval from stimulus onset, and the P3b amplitude was measured between 550–700 ms for target events in both groups of participants. Because distractors often did not elicit clear responses, the same intervals were used to measure the amplitudes of these responses, both for the numerals and syntactic violations.

#### 2.4.4. EEG source localization and functional connectivity analysis

For source localization, the EEG signal was segmented into epochs of 2048 ms duration. Epochs including target events, distractors, syntactic violations, or button presses were rejected from the analysis of functional connectivity. Epochs with an amplitude change of 100 μV or greater at any electrode were rejected. As a result, the analyzed dataset consisted of an average of 198 epochs per participant per condition (SD = 20.15). The Brainstorm toolbox (Tadel et al., 2011) was used to perform EEG source reconstruction, following the protocol of previous studies (Song et al., 2015; Huang et al., 2016; Pizzagalli., 2007). MNI’s Colin27 brain template was used to derive default anatomical regions, which, along with the default electrode locations, were entered into the forward boundary element head model (BEM) provided by the openMEEG algorithm (Gramfort et al, 2011).

For the modeling of time-varying source signals (current density) of all cortical voxels, a minimum norm estimate inverse solution was applied (sLORETA developed by Pascual-Marqui, 2002). The reconstructed dipole had a component perpendicular to the cortical surface. Averaging dipole strengths across voxels yielded mean current densities for 62 cortical areas, defined by the standardized parcellation scheme introduced by Klein & Tourville (2012). Finally, 18 left- and 18 right-hemispheric cortical regions of interest (ROIs) were selected for analysis. A region was defined as a ROI if it was linked with attentional, auditory, or speech processing in the literature (Saur et al., 2008, 2010). Table 2 contains the list of ROIs for the present study, together with their abbreviations. Source localization error has been estimated for each region by Toth and colleagues (2019). Reconstructed source activity was found to be ambiguous for only a few pairs of neighboring ROIs around Heschl’s gyrus, where reconstructed source activity could not be accurately distinguished from regions such as the supramarginal gyrus, insula, superior temporal and postcentral gyrus. Therefore, we do not draw conclusions from the FCs for these regions.

**Table 2.** EEG source regions and their abbreviations, grouped according to large-scale anatomical areas.

| Anatomical region | EEG source regions | Abbreviation |
| --- | --- | --- |
| **Frontal** | Caudal Middle Frontal Gyrus | MFGc |
|  | Rostral Middle Frontal Gyrus | MFGr |
|  | Orbitofrontal Gyrus | OFG |
|  | Inferior Frontal Gyrus | IFG |
|  | Superior Frontal Gyrus | SFG |
|  | Precentral Gyrus | PrCG |
| **Cingular** | Anterior Cingulate Cortex | ACC |
|  | Posterior Cingulate Cortex | PCC |
| **Temporal** | Fusiform Gyrus | FFG |
|  | Inferior Temporal Gyrus | ITG |
|  | Middle Temporal Gyrus | MTG |
|  | Superior Temporal Gyrus | STG |
| **Parietal** | Inferior Parietal Gyrus | IPG |
|  | Superior Parietal Gyrus | SPG |
|  | Supramarginal Gyrus | SMG |
|  | Paracentral Gyrus | PCG |
|  | Postcentral gyrus | PoCG |
|  | Precuneus | PCUN |

Functional connectivity (FC) was calculated by BrainWave (version 0.9.151.5), separately for each EEG epoch. FC was expressed as phase synchronization strength between each pair of ROIs, as measured by the phase lag index (PLI; Stam et al., 2007) in six frequency bands (delta: 0.5–4 Hz, theta: 4–8 Hz, lower alpha: 8–10 Hz, upper alpha: 10–12 Hz, beta: 13–30 Hz, gamma 30–80 Hz). PLI provides a value between 0 (random phase difference: minimum strength of FC) and 1 (constant phase difference: maximum strength of FC). The PLI calculation yielded symmetric matrices of 36*36 dimensions containing the PLIs for each pair of ROIs, separately for each frequency band. FC matrices were then averaged separately for each participant, condition, and frequency band, collapsed across stimulus blocks.

### 2.5. Statistical analysis

Statistical analyses of the behavioral and ERP data were performed by STATISTICA 13.1. For the hit rate (HR), distractor effect, d’, false alarm (FA) rate, and memory performance, separate repeated-measures ANOVAs were conducted for testing the effects of the within-group variable of PROSODY (intact vs. synthetically flattened vs. naturally suppressed prosody) and the between-group variable of GROUP (attended stream manipulated vs. ignored stream manipulated).

Amplitudes measured from Pz (as it showed the largest response for the components tested, see Supplementary Figure 1) for target numerals were compared using repeated-measures ANOVAs, with the factor of PROSODY, separately for the N2b and P3b components and for the two groups. The parietal ERP components elicited by the distractor numerals and syntactic violations were tested by one-tailed Student’s *t*-tests against 0 with FDR (false discovery rate) correction applied to account for multiple comparisons. The alpha level was set to 0.05. All significant results are reported. Sphericity violations were tested for all comparisons, and Greenhouse-Geisser correction was applied as necessary (the ε correction factor is then reported). The partial eta squared (ηp^2^) measure of effect size is also reported. Post-hoc tests were computed by Tukey’s HSD pairwise comparisons.

Statistical analysis of FC data was conducted by the network-based statistic (NBS) toolbox (Zalesky, Fornito & Bullmore, 2010). This algorithm first performs F tests on each functional connection of the given ROI pairs (N = 630 edges [(36 × 35)/2]) and collects the connections above the F threshold. Supra-threshold connection sets are created for the range of F thresholds between 3–10. The algorithm then searches for distinct subnetworks within these sets. Subnetworks are defined as fully connected networks in which any pair of nodes is connected by at least one path (i.e., each node has at least one connection). A family-wise error corrected p-value is obtained for each subnetwork using permutation-based mass univariate testing (i.e., testing is done by subnetworks rather than individual connections). During permutation testing, 10000 random subnetworks are created by repeatedly permuting the statistically tested condition’s FC value vectors, separately within each participant. The size (number of connections) of the largest supra-threshold subnetwork extracted from each permutation forms the distribution against which the supra-threshold subnetworks are tested (separately for each contrast). Networks with a size falling into the highest 5% of the distribution were regarded as significant. The significant subnetworks with the highest threshold within the F range were then selected for further network construction, setting the final threshold by determining the maximum F value that still resulted in at least one significant subnetwork for the given contrast, separately for the six EEG frequency bands. Next, within each subnetwork, the connections were ordered according to the size of the FC strength difference between the contrasted conditions. A maximum of 32 connections with the highest difference for each contrast (5% of the total number of connections) were submitted to post-hoc pairwise *t*-tests. Only edges with a significant (α = 0.05) difference were selected for interpretation as these characterize the largest FC strength difference contributing to the distinction tested by the contrast.

The effects of **PROSODY** were tested by pairwise contrasts, separately for the two groups and the EEG bands: 1) intact prosody vs. synthetically flattened prosody and 2) intact prosody vs. naturally flattened prosody. The BrainNet Viewer toolbox (Xia & Wang, 2013) was used to visualize the FC analysis results over the cortical surface applying the BrainMesh_ICBM152 surface template to the nodes representing ROIs (locations represented by the standard MNI coordinates).

## 3. Results

### 3.1. Behavioral responses

Figure 2 shows the effects of PROSODY and GROUP on HR (A), d’prime (B), the distractor effect (C), FA rate (D), and recognition memory performance (E). Table 3 contains the F, p, and ηp^2^ values of these measures yielded by the ANOVA test. While both GROUP and PROSODY had a significant main effect on all but recognition memory performance, these were caused by the significant interaction between these factors. The post-hoc tests showed that listeners in the attended stream manipulated group responded differently to naturally suppressed prosody than to any condition in either group: HR and d’ were lower, the distractor effect and FA rate higher for this than for any other combination of condition and group (p < 0.001, all). Recognition memory performance was lower for naturally suppressed, as well as for synthetically flattened prosody compared to the intact speech condition, with no significant GROUP effect or interaction.

**Figure 2.**
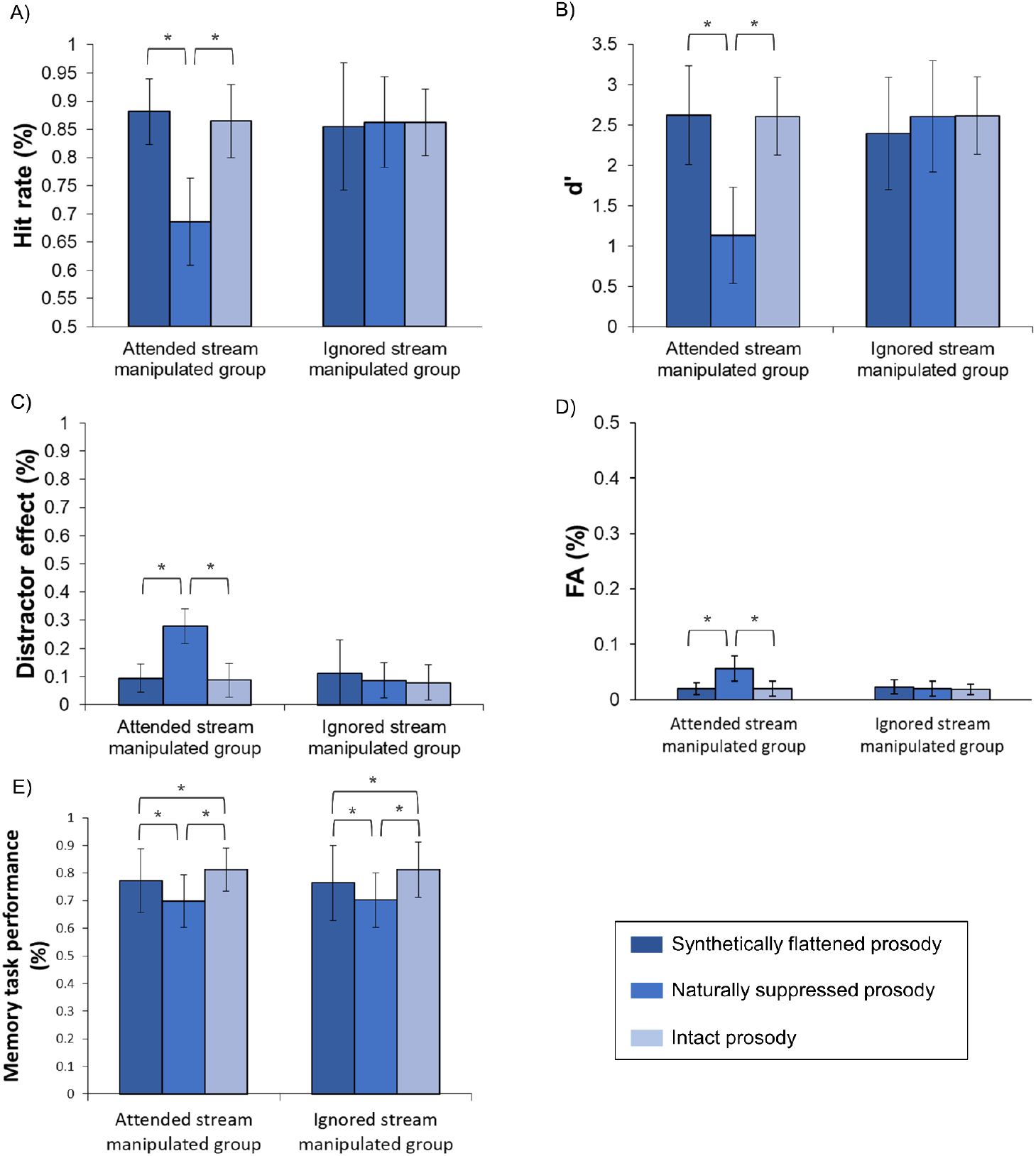
Effects of PROSODY and GROUP on HR (A), d’ (B), the distractor effect (C), FA rate (D), and recognition memory performance (E).

**Table 3.** F, p, and ηp^2^ values of the statistically significant behavioral effects. Where sphericity was violated, Greenhouse-Geisser ε is reported.

| Variable | GROUP | PROSODY | GROUP * PROSODY |
| --- | --- | --- | --- |
| HR | $F(1, 48) = 7.22$<br>$p < 0.01$<br>$\eta p^2 = 0.13$ | $F(2, 96) = 46.25$<br>$\varepsilon = 0.67$<br>$p < 0.001$<br>$\eta p^2 = 0.49$ | $F(2, 96) = 51.07$<br>$\varepsilon = 0.67$<br>$p < 0.001$<br>$\eta p^2 = 0.51$ |
| $d'$ | $F(1, 48) = 8.30$<br>$p = 0.01$<br>$\eta p^2 = 0.15$ | $F(2, 96) = 49.67$<br>$\varepsilon = 0.72$<br>$p < 0.001$<br>$\eta p^2 = 0.51$ | $F(2, 96) = 54.54$<br>$\varepsilon = 0.72$<br>$p < 0.001$<br>$\eta p^2 = 0.53$ |
| Distractor effect | $F(1, 48) = 13.89$<br>$p < 0.001$<br>$\eta p^2 = 0.22$ | $F(2, 96) = 58.20$<br>$\varepsilon = 0.65$<br>$p < 0.001$<br>$\eta p^2 = 0.55$ | $F(2, 96) = 58.20$<br>$\varepsilon = 0.65$<br>$p < 0.001$<br>$\eta p^2 = 0.58$ |
| FA | $F(1, 48) = 11.66$<br>$p = 0.001$<br>$\eta p^2 = 0.20$ | $F(2, 96) = 60.37$<br>$\varepsilon = 0.69$<br>$p < 0.001$<br>$\eta p^2 = 0.56$ | $F(2, 96) = 63.11$<br>$\varepsilon = 0.69$<br>$p < 0.001$<br>$\eta p^2 = 0.57$ |
| Recognition memory | | $F(2, 96) = 22.48$<br>$p < 0.001$<br>$\eta p^2 = 0.32$ | |

### 3.2. ERP responses

In the attended stream manipulated group (Figure 3, left panel), a significant main effect of PROSODY was observed for the target N2b amplitudes (F(2,48) = 8.056; ε = 0.671; p < 0.01; ηp^2^ = 0.251). The post-hoc test revealed that N2b amplitudes in the intact speech condition were significantly lower than in the synthetically flattened or the naturally suppressed prosody conditions (p < 0.01, both), while the latter two did not significantly differ from each other (p = 0.763). In contrast, the main effect of PROSODY did not reach significance for the P3b amplitudes (p = 0.067); the tendency appears to stem from numerically lower P3b amplitudes in the naturally suppressed prosody than in the other two conditions. ERPs measured for distractor numerals and syntactic violations in the ignored stream did not reach significance after FDR correction (p > 0.227, all), except for syntactic violations appearing in the to-be-ignored stream in the naturally suppressed prosody condition (p < 0.05), for which a response significantly different from zero was observed in the P3 latency range. Note, however, that this response was negative-going, continued beyond the P3 latency range, and showed negative amplitude values (see Supplementary Figure 2). Thus it may rather be an N400 type of response.

**Figure 3.**
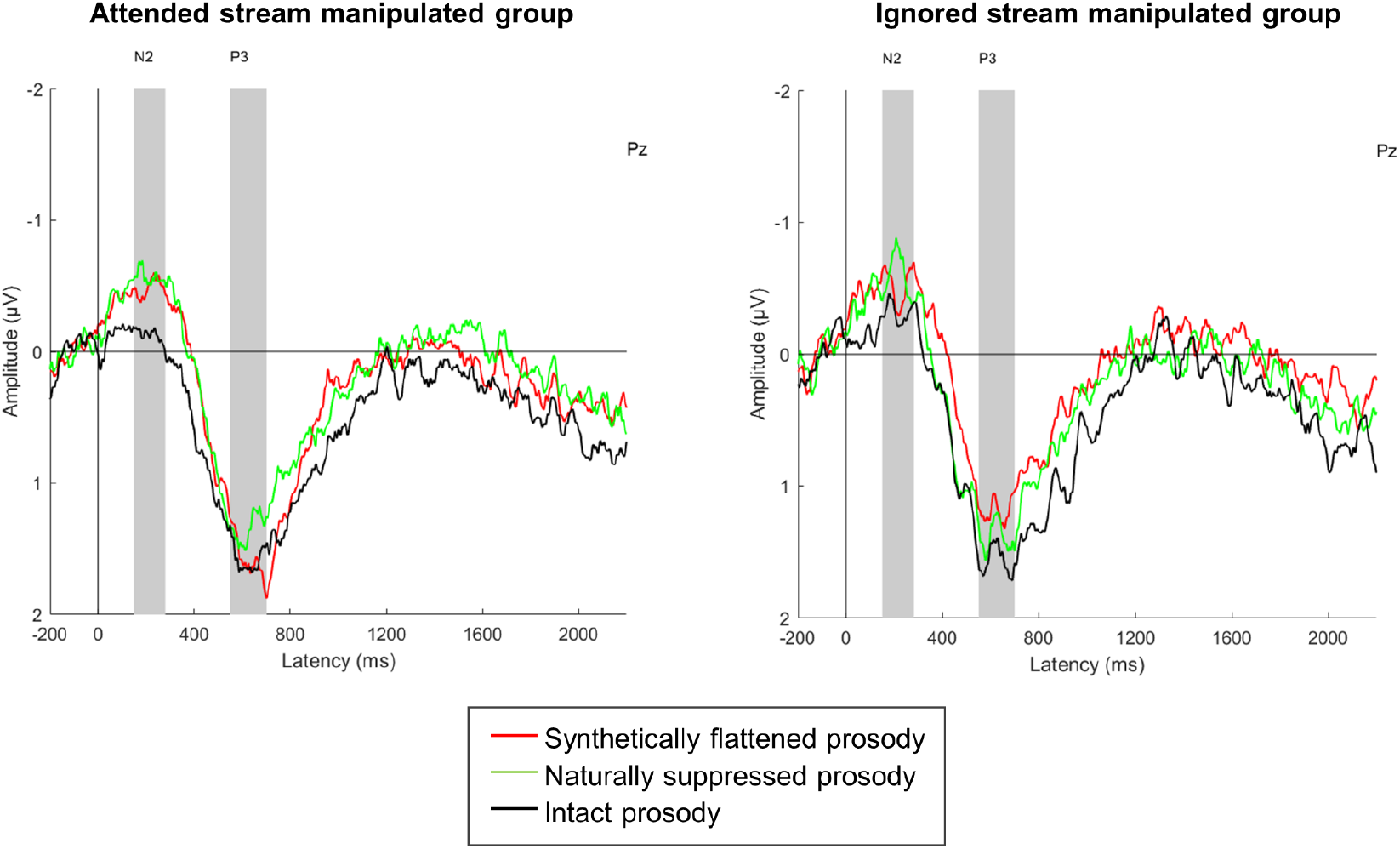
Group-averaged (N = 25) target-related parietal (Pz) ERPs elicited in the attended (left) and the ignored stream manipulated group (right). Zero latency is at the onset of the numeral word. The measurement windows for N2b and P3b are shown by gray vertical bands. In the attended stream manipulated group, target N2b amplitude in the intact prosody condition was significantly lower than in the other conditions (left panel, black line).

In the ignored stream manipulated group, a significant PROSODY main effect was found for the target P3b (F(2,48) = 4.373; ε = 0.769 p < 0.05; ηp^2^ = 0.154) but not for the N2b amplitudes (p = 0.150; Figure 3, right panel). This result was due to the significantly larger P3b elicited by targets in the intact speech compared in the synthetically flattened speech condition (p < 0.05). No further significant differences were found in the post-hoc test (p > 0.281). Neither the distractor numerals nor the syntactic violations elicited significant ERP components (p > 0.169, all; Supplementary Figure 2).

### 3.3. EEG Functional connectivity

Networks significantly influenced by the prosody manipulation were identified in the delta, theta, alpha, and gamma bands. Supplementary Table 1 shows the number of connections (node degree) within the networks showing significant effects, separately for each brain region (ROI) and frequency band. Nodes with the highest number of connections can be regarded as hubs of interregional communication. The networks characteristically differ from each other by hub locations and the relative contribution of the different lobes. Significant networks are illustrated on pairs of plots of the cortical surface (shown by lateral views of the two hemispheres) with the hub regions marked with their corresponding label (Figure 4).

**Figure 4.**
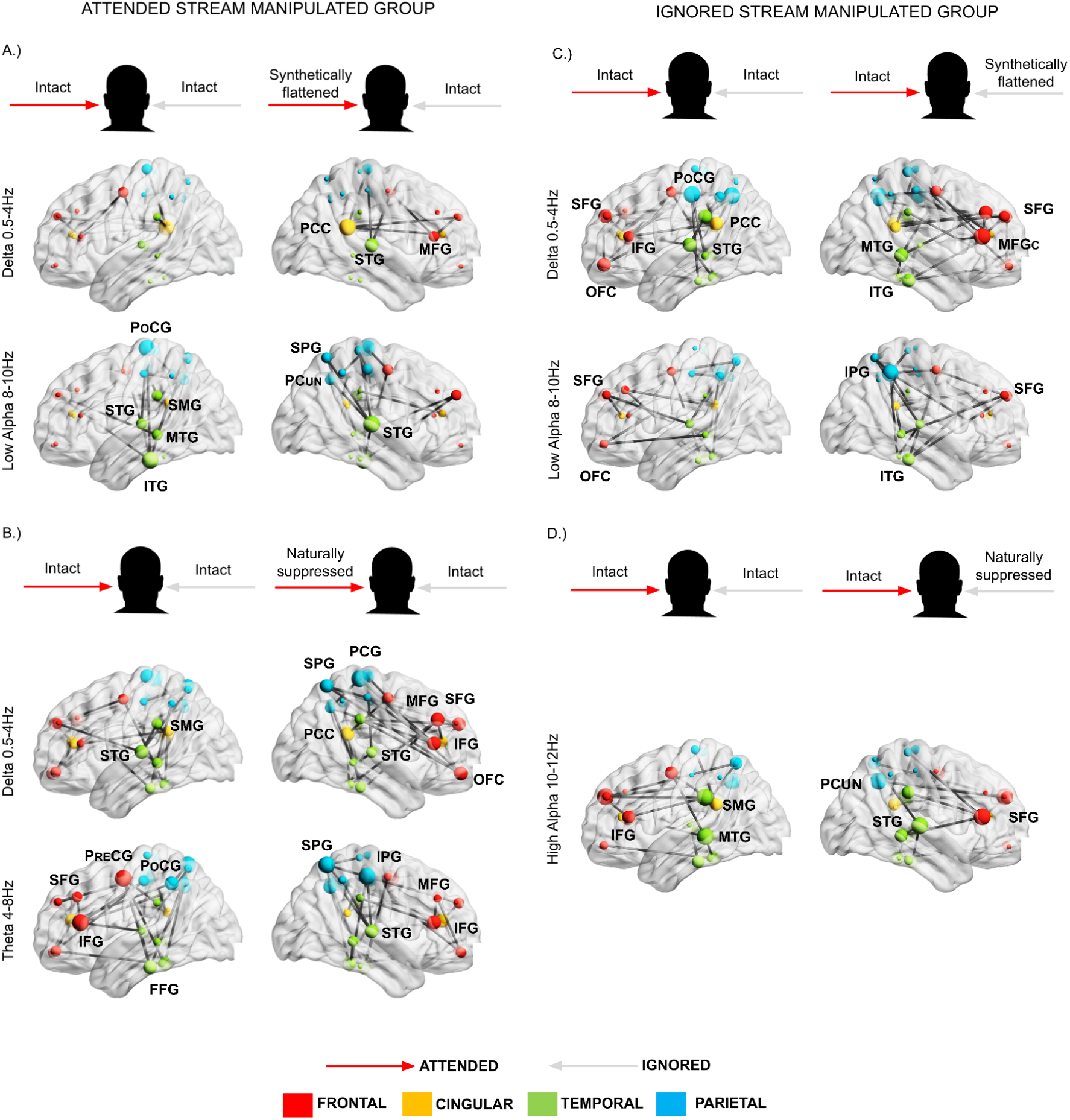
Functional brain networks significantly affected by PROSODY in the attended stream manipulated (panels A and B, left column) and the ignored stream manipulated group (panels C and D, right column). Brain networks are plotted on the left and right lateral view of the cortical surface. Dots represent the spatial locations of the ROI’s quasi center (nodes) defined in MNI space. The size of the node represents the number of connections of the node within the network. The color of the node indicates the cortical lobe: red – frontal; yellow – cingular; green – temporal; blue–parietal cortex. The two compared conditions are shown on the top of each panel. Red arrows mark the to-be-attended, gray the to-be-ignored stream.

The PROSODY manipulation resulted in the activation of different brain networks in the two groups (Fig. 4). In the attended stream manipulated group, a network (9 connections, 10 nodes) operating in the delta band (0.5–4 Hz) showed significantly different synchronization strength between the intact and the synthetically flattened speech conditions (Figure 4A, upper panel). This network is dominant in the right hemisphere, with the main hub nodes in PCC and STG. Another network (18 connections, 17 nodes) sensitive to the same contrast was observed in the low alpha band (8–10 Hz; Figure 4A, lower panel). This network is bilateral, with connections within the parietal and temporal lobes, and between parietal, temporal, and frontal regions. Its hub nodes are in the left ITG, PoCG and MTG, as well as in the right STG, PCUN and SPG.

In the same group, comparison between the naturally suppressed prosody and the intact speech condition yielded significant networks in the delta (0.5–4 Hz; 33 connections, 32 nodes; Figure 4B, upper panel) and the theta band (4–8 Hz; 33 connections, 32 nodes; Figure 4B, lower panel). Both of these networks involve frontal, temporal, and parietal regions. The delta network has a large number of hubs, including the right frontal lobe (OFC, IFG, MFG, SFG), right parietal areas (SPG, PCG), as well as STG, PCC and SMG bilaterally. Nodes with the highest number of connections in the theta network are located bilaterally in the parietal (PoCG, SPG, IPG) and frontal cortex (PreCG, IFG, SFG), in the left FFG, and the right SPG, IPG, and STG.

In the ignored stream manipulated group, networks sensitive to the contrast between the intact and the synthetically flattened speech were observed in the delta (0.5–4 Hz; 33 connections, 31 nodes; Figure 4C, upper panel) and low alpha band (8–10 Hz; 24 connections, 23 nodes; Figure 4C, lower panel). The delta-band network bilaterally connects frontal, temporal, and parietal areas. Hub nodes reside in the temporal (MTG, ITG, STG) parietal (POCG), cingulate (PCC), and frontal (OFC, IFG, MFG, SFG) regions. The low-alpha-band network involves hub areas in the left SFG and OFC, as well as the right IPG, SFG, and ITG.

Contrasting intact speech and naturally suppressed prosody in the same group, a significant network operating in the high alpha band was found (10–13 Hz; 21 connections, 21 nodes; Figure 4D) with predominantly fronto-parietal and fronto-temporal connections. Hubs are located in frontal (SFG, IFG), parietal (PCUN), and temporal (MTG, STG, SMG) areas.

## 4. Discussion

We assessed the influence of prosody on segregating and selecting one speech stream from two concurrent speech streams. Prosody was separately manipulated for the to-be-selected and the to-be-ignored speech stream. We expected that speech with degraded prosody would make stream/target selection more difficult, but affect stream segregation to a lesser degree (Hypothesis 1) and that the synthetic manipulation applied in this study (flattening the dynamic of F0 in continuous speech) would have a larger effect than when the speakers themselves try to suppress prosody while voicing the articles (Hypothesis 2). Our results indicate that a natural, intact prosody does indeed facilitate attentional processes, whereas it is not a similarly effective cue of stream segregation: behavioral effects were significantly stronger when speech with degraded prosody was to be attended than when it was to be ignored. However, we found that prosody suppressed by the speakers had a far larger effect than flattening the F0, especially when it was applied to the attended speech stream. In the following, we will discuss the behavioral and neurophysiological (ERP and FC) findings in more detail.

### 4.1. Effects of degraded prosody on solving the cocktail-party problem

#### 4.1.1. Behavioral results

Prosodically degrading the attended stream induced lower HR and higher distraction, whereas similar manipulation of the ignored stream did not result in significant target detection effects. This suggests that stream segregation or selection was more difficult with degraded prosody. Because the effect was not present when the manipulated stream was ignored, the effect cannot stem from stream segregation alone. In addition to degraded stream selection, a prosody effect on target selection is also possible, although that alone would not explain the observed increase in distraction.

When naturally prosody-suppressed speech was to be attended, the FA rate also increased. Further, recognition memory performance was similarly affected when prosody was manipulated in the attended or the ignored stream: lower for naturally suppressed prosody than for intact or synthetically flattened prosody. Increases in false alarm rate can be either attributed to distraction from the ignored stream and/or more difficult target detection, while memory performance may have decreased due to difficulties in telling the contents of the two streams apart. Especially the latter effect suggests problems with auditory stream segregation. Therefore, degradation of prosody does affect stream segregation to an extent, albeit this is a weaker effect than the effect of prosody on selective attentional processes, since participants successfully compensated for it in their behavioral performance.

#### 4.1.2. ERP results

N2b amplitudes were higher when the prosody of the attended stream was manipulated in any way, than in the case of an intact attended stream. Since N2b amplitudes increase with greater processing demands (Patel & Azzam, 2005), this result implies that the detection of targets in prosodically manipulated speech imposes an increased cognitive load. The statistically significant response (possibly N400 – as was also found in Szalárdy et al., 2018) to syntactic violations in the naturally suppressed prosody condition suggests that attention may have switched from the attended to the ignored speech stream, at least in the naturally suppressed prosody condition. Together with the behavioral data, these results suggest that suppressed prosody makes stream selection more difficult.

Target P3b amplitudes were lowest in the naturally suppressed prosody condition. Since, according to Polich (2007), smaller P3b amplitudes are associated with more difficult target detection tasks, where more attentional resources are required, this result implies that degraded prosody may also affect target detection. As the P3b amplitude effect did not reach significance (it was only a tendency), this effect may be smaller than the effect on stream selection.

In the ignored stream manipulated group, the P3b amplitudes elicited by targets (i.e. numerals in the attended stream, which was always intact in this group) were larger when the ignored speech was intact than when the ignored speech was synthetically flattened. Together with the effect of ignored naturally suppressed prosody on recognition performance, this result supports our suggestion that prosody may also affect auditory stream segregation, although to a lesser degree than stream selection.

#### 4.1.3. Functional connectivity results

Slow oscillations (1–7 Hz) support the processing of speech in multi-talker settings by tracking the attended speech stream (Golumbic et al., 2013). Therefore, if it is stream selection rather than stream segregation which is influenced by prosody, one should expect that functional networks in the delta (0.5–4 Hz) and theta band (4–8 Hz) will be sensitive to the current manipulations. In accordance with Golumbic and colleagues’ (2013) study, the results show that FC differences between the intact and synthetically flattened conditions, whether attended or ignored, emerged in networks in the delta band. Delta-band differences were supplemented by differences in theta band in the naturally suppressed prosody speech condition in the attended stream manipulated group. Slow oscillatory networks were especially widely distributed upon attending to naturally prosody-suppressed speech, with frontal (MFG, SFG, OFC), parietal (PCG, SPG) and temporal (STG) hubs. The involvement of the frontoparietal attention network (Corbetta et al., 2002) in speech perception, and its coupling to the STG – an important area for speech processing (Hickok & Poeppel, 2007; Vander Ghinst et al., 2016) – indicates that more top-down control was needed in order to follow the to-be-attended speech stream effectively when prosody was highly atypical (cf. Park et al., 2015; Tóth et al., 2019). Further, the theta band oscillatory network showing significant coupling in this condition included the right IFG and right superior temporal areas, which have been previously shown to be involved in prosody processing (Sammler et al., 2015). The right IFG and STG were also observed as hubs of a high alpha band network while ignoring speech with naturally suppressed prosody. This finding is in line with the expected suppression effect on these speech processing areas. It is worth noting that a widely distributed delta (0.5–4 Hz) band network was activated when the participants were following intact speech while ignoring synthetically flattened speech: we suggest that top-down regulation of the regions in the ventral temporal cortex involved in speech recognition (STG, MTG, ITG; Hickok & Poeppel, 2007) can be observed also in this case, complementing the results found for the P3b ERP component.

Besides the observed sensitivity of slow oscillations, significant connectivity differences were found in the low and high alpha frequency bands (8–10 Hz and 10–12 Hz, respectively). Alpha oscillations have been implicated in the inhibition of background noise during speech processing (Obleser et al., 2007; Strauß et al., 2014; Tóth et al., 2019). Accordingly, the current data show differences in alpha coupling between conditions where our behavioral results suggest difficulties in inhibiting the distracting speech stream (conditions with a high distractor effect) and conditions where the participants managed to suppress distractors better. In the naturally suppressed prosody condition of the attended stream manipulated group, where the distractor effect was highest, no functional network was observed in the alpha frequency band. In the other conditions, where distractor effect was low, functional connectivity was significant in either the low- or high-alpha band. The topography of these networks indicates that the temporal areas involved in speech processing (SMG, STG, MTG, ITG; Hickok & Poeppel, 2007) received top-down inhibition from parietal (PoCG, SPG, IPG, PCUN) and/or frontal areas (SFG) involved in attentional processes (Corbetta et al., 2002; Bahmani et al., 2019).

In sum, the FC network findings are fully compatible with and complement the effects found on behavioral and ERP measures. They show that the speech- and specifically the prosody-processing networks involved in selecting the to-be-attended stream and suppressing the to-be-ignored stream have been sensitive to the current prosody manipulations. While some of the similarities between the networks found in the attended stream and the ignored stream manipulated groups may stem from effects on auditory stream segregation, differences between them probably represent that attending and ignoring the same manipulated speech poses different difficulties to the auditory sensory-cognitive system (cf. in the next section).

### 4.2. The roles of different prosodic cues in solving the cocktail-party problem

We also hypothesized that completely averaging out the F0 of speech would have a more detrimental effect on successfully solving the cocktail party problem than naturally induced prosody suppression, since in the latter case, pitch is more variable, and F0 changes are a prominent aspect of speech prosody (Meyer, 2018; see also the current analysis in Section 2.2). However, this hypothesis did not hold: naturally suppressing speech prosody (monotonous speech, i.e. little variation in F0) proved to be more difficult to follow than synthetically F0-flattened speech (i.e. no variation in F0). This suggests that intonation is not the only prosodic cue which aids listeners in segregating and selecting speech streams.

Although there might be other prosodic differences between the two manipulations, we showed that there was a difference in speech rate (see Table 1): natural suppression of prosody yielded faster speech, possibly with shorter pause intervals, which likely hindered the parsing of speech into meaningful units, interfering with speech comprehension, and thus rendering the task of segregating and selecting a speech stream, as well as detecting targets within it more difficult. Lower memory performance compared to synthetically flattened speech, irrespective of whether speech with natural prosody suppression was attended to or ignored, supports the notion of the effect on stream segregation. Higher distraction and the elicitation of N400 by syntactic violation in the unattended stream when the manipulated speech was attended indicate degraded stream selection, with the lack of significant low-frequency networks sensitive to the difference from natural speech in the ignored stream being compatible with this inference. Difficulties in target detection are reflected in the higher FA rate. In a study using similar stimuli and task, Szalárdy and colleagues (2020) found significantly lower hit rates and memory performance and significantly higher N2b amplitudes, when both streams consisted of speech segments artificially accelerated by 10% (the approximate effect of “natural prosody suppression” in the current study) than when intact speech was delivered in both streams. These results are fully compatible with the current ones. On the other hand, in the comparable conditions of Szalárdy and colleagues (2020), no significant distractor effect was found and no 400 was elicited by syntactic violations embedded in the to-be-ignored stream. These differences might either stem from the fact that only one of the streams was manipulated in the current study (while both streams have been manipulated in parallel in Szalárdy et al., 2020), and/or because natural suppression of prosody does not only affect speech rate, but also other cues of prosody. Thus, speech rate is an important suprasegmental feature aiding speech processing in multi-talker situations.

In addition to the effects of speech rate, F0 also plays a role in solving the cocktail party problem. Lower P3b amplitudes were obtained when synthetically flattened than when intact speech was to be ignored, while a P3b with numerically higher amplitude was elicited for synthetically flattened speech than for naturally suppressed speech, when they were attended. This asymmetry suggests that ignoring synthetically flattened speech may be easy, while when listening to it, target detection may pose some problems. In sum, prosody affects how well listeners can solve the cocktail party problem. Here we found significant effects of speech rate and F0, while future studies may test the effects of other cues of prosody.

### 4.3. Summary

The current results indicate that prosody supports stream selection, auditory stream segregation, and target detection, albeit to a different degree. While a strong effect on stream selection was found, the effect on auditory stream segregation was lower, and that on target detection only marginal. The underlying brain networks mainly operate on the lower (delta, theta) frequency bands, indicating top-down control over sensory and integrative processes of speech perception, as well as in the alpha band. The latter are likely involved in suppressing the ignored speech. Speech rate (probably including pause intervals) appears to be a strong cue supporting the sensory/cognitive system in solving the cocktail-party problem, while the trajectory of the fundamental frequency has a weaker effect on it.

**Data is available at https://osf.io/68auf/**.

## Author Notes

This work was funded by the Hungarian National Research Development and Innovation Office (OTKA project PD123790 to TB and K132642 to IW) and by the Hungarian Academy of Sciences (LP2012-36/2012 to IW and the János Bolyai grant awarded to TB). The authors are grateful to Ágnes Palotás, Gábor Orosz, and László Hunyadi for text editing, to Ferenc Elek and Péter Scherer for voicing the articles, to László Liszkai for audio recording and editing, to Botond Hajdu for implementing the experimental design, to Zsuzsanna Kovács for data collection, and to Bálint File for programming the EEG source reconstruction algorithm.

The authors declare no conflict of interest.

