## Supplementary material for "Speech prosody supports speaker selection and auditory stream segregation in a multi-talker situation"

**Supplementary Figure 1.** Scalp distributions for the N2 and P3 ERP components elicited by *target numerals.*

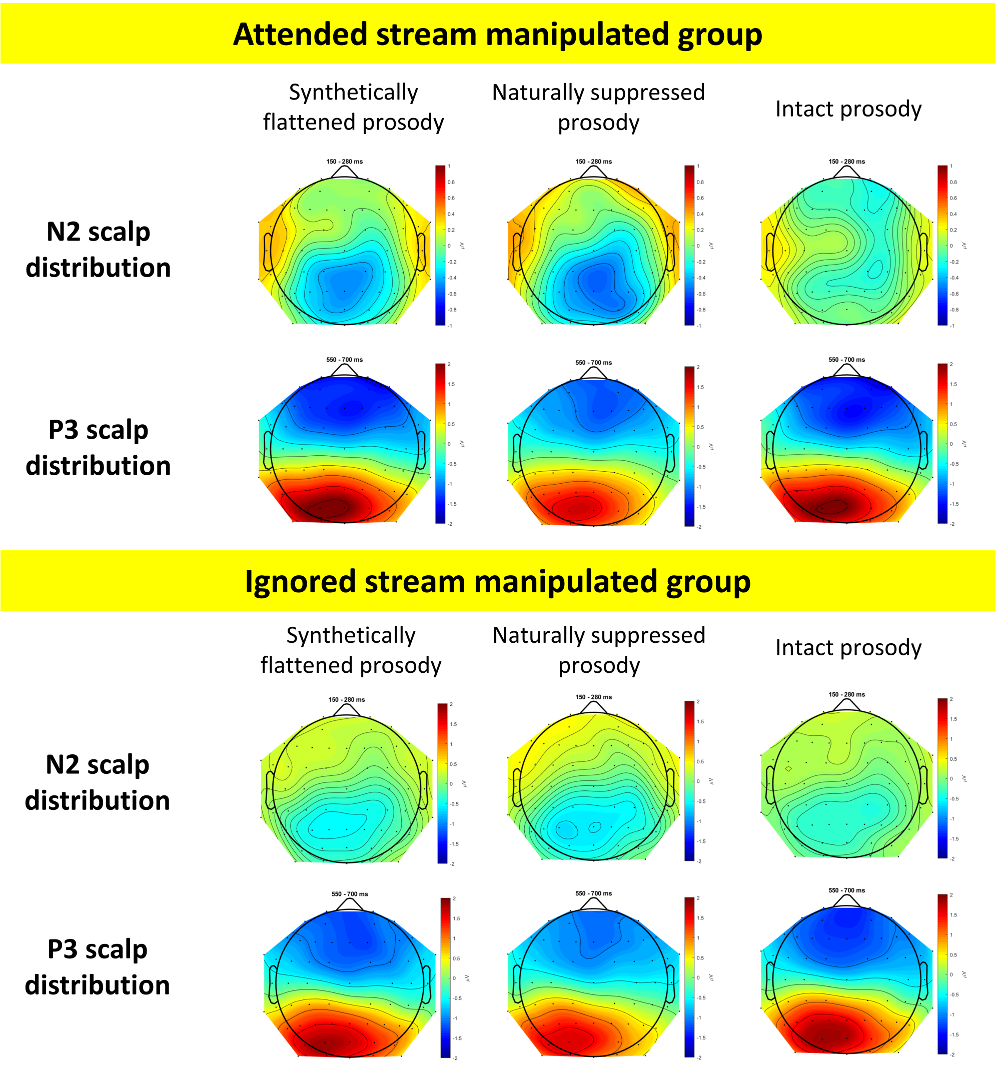

**Supplementary Figure 2.** The N2 and P3 ERP components elicited by the distractor numerals were non-significant in all groups and conditions. Results were the same for the distractor syntactic violations, except for the response in the P3 latency range, which significantly differed from zero ($p < 0.05$) in the naturally suppressed prosody condition in the attended stream manipulated group.

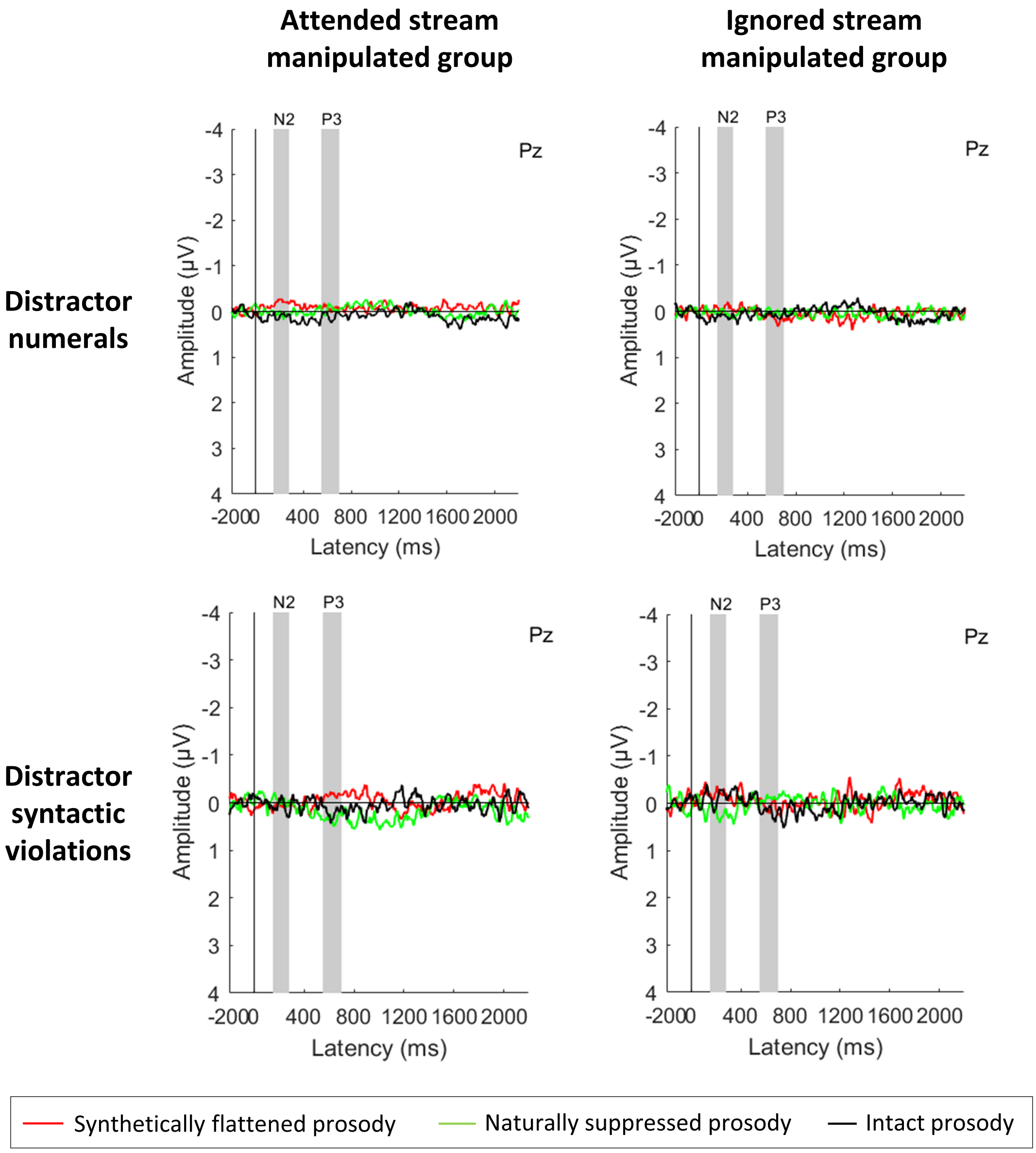

**Supplementary Table 1.** The number of connections within the subnetworks showing significant connectivity in the ATTENDED and IGNORED STREAM MANIPULATED GROUP, respectively.

| | **ATTENDED STREAM MANIPULATED GROUP** | | | | **IGNORED STREAM MANIPULATED GROUP** | | | |
|---|---|---|---|---|---|---|---|---|
| | Intact vs Synthetically flattened | | Intact vs Naturally suppressed | | Intact vs Synthetically flattened | | | Intact vs Naturally suppressed |
| | Delta | Low Alpha | Delta | Theta | Delta | Low Alpha | Gamma | High Alpha |
| **'ACC' L** | 0 | 0 | 2 | 0 | 1 | 1 | 0 | 1 |
| **'ACC' R** | 1 | 1 | 2 | 3 | 1 | 0 | 0 | 0 |
| **'MFGC'L** | 0 | 0 | 0 | 2 | 0 | 2 | 0 | 0 |
| **'MFGC'R** | 0 | 0 | 3 | 1 | 3 | 0 | 0 | 0 |
| **'FFG'** | 0 | 0 | 2 | 3 | 2 | 2 | 0 | 2 |
| **'FFG'** | 0 | 0 | 1 | 2 | 1 | 0 | 0 | 0 |
| **'IPG'** | 0 | 0 | 1 | 3 | 2 | 2 | 0 | 0 |
| **'IPG'** | 0 | 1 | 0 | 1 | 0 | 6 | 0 | 0 |
| **'ITG'** | 0 | 4 | 2 | 3 | 1 | 1 | 0 | 2 |
| **'ITG'** | 0 | 0 | 2 | 0 | 3 | 4 | 0 | 1 |
| **'PCC'** | 0 | 1 | 2 | 0 | 3 | 0 | 0 | 2 |
| **'PCC'** | 4 | 0 | 3 | 1 | 2 | 2 | 0 | 0 |
| **'OFC'** | 0 | 0 | 2 | 2 | 3 | 2 | 0 | 1 |
| **'OFC'** | 0 | 2 | 3 | 2 | 1 | 0 | 2 | 0 |
| **'MTG'** | 0 | 2 | 2 | 1 | 1 | 1 | 0 | 3 |
| **'MTG'** | 0 | 0 | 1 | 2 | 3 | 2 | 0 | 2 |
| **'PCG'** | 2 | 4 | 3 | 1 | 1 | 0 | 0 | 1 |
| **'PCG'** | 0 | 2 | 4 | 1 | 2 | 1 | 2 | 1 |
| **'IFG'** | 0 | 1 | 1 | 4 | 2 | 0 | 0 | 2 |
| **'IFG'** | 2 | 0 | 3 | 3 | 4 | 0 | 0 | 3 |
| **'PoCG'** | 0 | 4 | 1 | 2 | 4 | 1 | 1 | 0 |
| **'PoCG'** | 0 | 2 | 1 | 4 | 1 | 0 | 0 | 0 |
| **'PrCG'** | 2 | 0 | 2 | 4 | 2 | 2 | 0 | 2 |

| | | | | | | | | |
|---|---|---|---|---|---|---|---|---|
| **'PRCG'** | 0 | 2 | 2 | 1 | 2 | 2 | 0 | 0 |
| **'PCUN'** | 0 | 1 | 1 | 2 | 4 | 0 | 0 | 0 |
| **'PCUN'** | 0 | 2 | 2 | 3 | 2 | 0 | 0 | 3 |
| **'MFGR'** | 0 | 0 | 0 | 1 | 1 | 2 | 0 | 1 |
| **'MFGR'** | 0 | 2 | 0 | 0 | 2 | 2 | 0 | 0 |
| **'SFG'** | 1 | 0 | 2 | 1 | 3 | 0 | 0 | 3 |
| **'SFG'** | 1 | 0 | 3 | 2 | 2 | 3 | 0 | 2 |
| **'SPG'** | 0 | 2 | 1 | 1 | 0 | 2 | 0 | 2 |
| **'SPG'** | 0 | 2 | 3 | 4 | 0 | 2 | 1 | 0 |
| **'STG'** | 1 | 0 | 3 | 1 | 3 | 2 | 0 | 0 |
| **'STG'** | 3 | 2 | 2 | 3 | 0 | 2 | 2 | 3 |
| **'SMG'** | 1 | 2 | 2 | 1 | 3 | 2 | 0 | 3 |
| **'SMG'** | 0 | 4 | 2 | 1 | 1 | 0 | 2 | 2 |
